# Naturalistic recall preserves temporal specificity despite cortical feature reorganization

**DOI:** 10.64898/2026.11.03.763513

**Authors:** Shikang Peng, Brian Levine, Bradley R. Buchsbaum

## Abstract

Although sensory input arrives in a largely continuous stream, recall is temporally compressed, discontinuous, and often structured as events. However, the temporal specificity of recall-related neural activity within events, and the separation and cortical organization of recalled features relative to perception, remain poorly characterized. We analyzed functional MRI data recorded during viewing and free recall of an episode of *Sherlock*. A data-driven alignment between viewing and recall data identified neural recapitulation of corresponding narrative events and revealed information that distinguished moments within events, particularly in the association cortex. Features ranging from low-level visual to semantic information predicted recall reinstatement. However, differences in predictive performance among feature levels became smaller during recall, and their organization along the cortical sensory-to-association gradient reversed relative to viewing. Thus, temporal specificity within events can be preserved despite substantial reorganization of feature representations across the cortex during recall.

## Introduction

In the BBC series *Sherlock*, a crime scene investigation centers on the mysterious death of a woman dressed in pink. The scene contains multiple levels of visual information, from color and spatial layout to objects, faces, and scene meanings, whose relevance can change as the narrative unfolds. When this episode is later recounted to a friend, the features and sequential contents embedded in this perceptual stream are reconstructed in the absence of sensory input present during viewing. In this study, we investigated the extent to which the neural organization of the originally perceived information is retained during free recall.

Extensive evidence from neuroimaging studies using cued recall shows that retrieval reinstates perceptual content, including stimulus categories and low-level visual features in early sensory cortex^1,2,3^. While such findings establish a match between information expressed during perception and retrieval, there are also systematic perception-to-memory transformations in both where stimulus information is expressed and how it is represented. Stimulus information prominent in the occipitotemporal cortex during perception is later expressed in the frontoparietal cortex during retrieval^4,5,6^, and representations retained within the visual cortex exhibit different spatial tuning properties during perception and memory^7^. Thus, perceptual information can be reinstated even as its neural organization is transformed.

While such studies are informative, perception in naturalistic events involves continuously evolving features spanning multiple levels of representation. Neuroimaging studies of films or stories have shown that neural responses are organized across temporal scales, with early sensory cortex tracking rapidly changing input and higher-order regions integrating information over increasingly extended contexts^8,9^. During memory retrieval, event-specific patterns elicited during movie viewing are reinstated^10,11^, and data-driven event models recover corresponding event structure across viewing and recall^12^. Reinstatement of short movie segments also varies with the temporal precision of spoken recollections^13^. However, these findings do not establish whether recall activity distinguishes different moments within the same event, beyond identifying the event being remembered. This distinction matters because reinstating an event-level representation need not imply that its constituent contents are reconstructed with moment-specific variation. Moreover, even when such information is retained, its feature-level organization may differ from that established during perception. Characterizing both properties within the same framework is therefore important for understanding how evolving experiences are represented during naturalistic recall.

Convolutional neural networks (CNNs) provide a framework for parameterizing multi-levels of information in naturalistic visual input. Features derived from CNN layers systematically predict neural activity across the human visual hierarchy^14,15^. These representations have also characterized visual imagery^16^ and feature-specific reactivation during cued recall of individual images^3^, where participants deliberately reconstruct a specified visual target. To study how feature organization happens in a continuously evolving naturalistic stream involving self-guided selection and verbal reconstruction, we need an alignment approach that maps recall contents with the corresponding moments in the movie timeline.

Here, we resolved feature content within events to characterize how neural representations change from movie viewing to free recall. We analyzed fMRI data acquired while participants watched an extended movie and subsequently recounted it^10^. We developed a data-driven framework based on optimal transport^17^ to map each recall moment onto a weighted combination of viewing moments, accommodating temporal compression and reconstructive integration^18^. We then trained feature-specific encoding models on CNN-derived movie representations to predict neural activity. We found that temporally-resolved representations reliably predicted neural activity and produced a significant improvement in prediction performance over event-level representations. At the feature levels, the neural organization present during viewing converged markedly during recall – an effect that cannot be explained by weaker model prediction or temporal smoothing induced by the alignment. Naturalistic recall therefore preserved evolving structure within events while transforming the differentiated feature hierarchy that organized perception.

## Results

The fMRI data used in this study involved 16 participants who watched a 48-min movie and subsequently recalled it freely during scanning^10^. To study feature representations from encoding to recall, we trained feature-specific encoding models on each cortical parcel (300-parcel)^19^ using four levels of movie features (low-, mid-, high-level, and semantic) derived from VGG-16^20^ to predict neural activity during viewing. Using encoding movie annotations and participants’ recall transcripts^10,21^ (Fig. 1a-b), we created a data-driven framework based on fused optimal transport (OT)^17^ to align each recall TR to a weighted distribution over encoding TRs, which jointly considers semantic correspondence between these states (*what* was recalled; Wasserstein term) and relational organization of features (*when* was recalled; Gromov term; Fig. 1c). The resulting transport plans were validated against independent manual scene annotations and used to reconstruct recall features to predict feature reactivation during recall.

**Figure 1.**
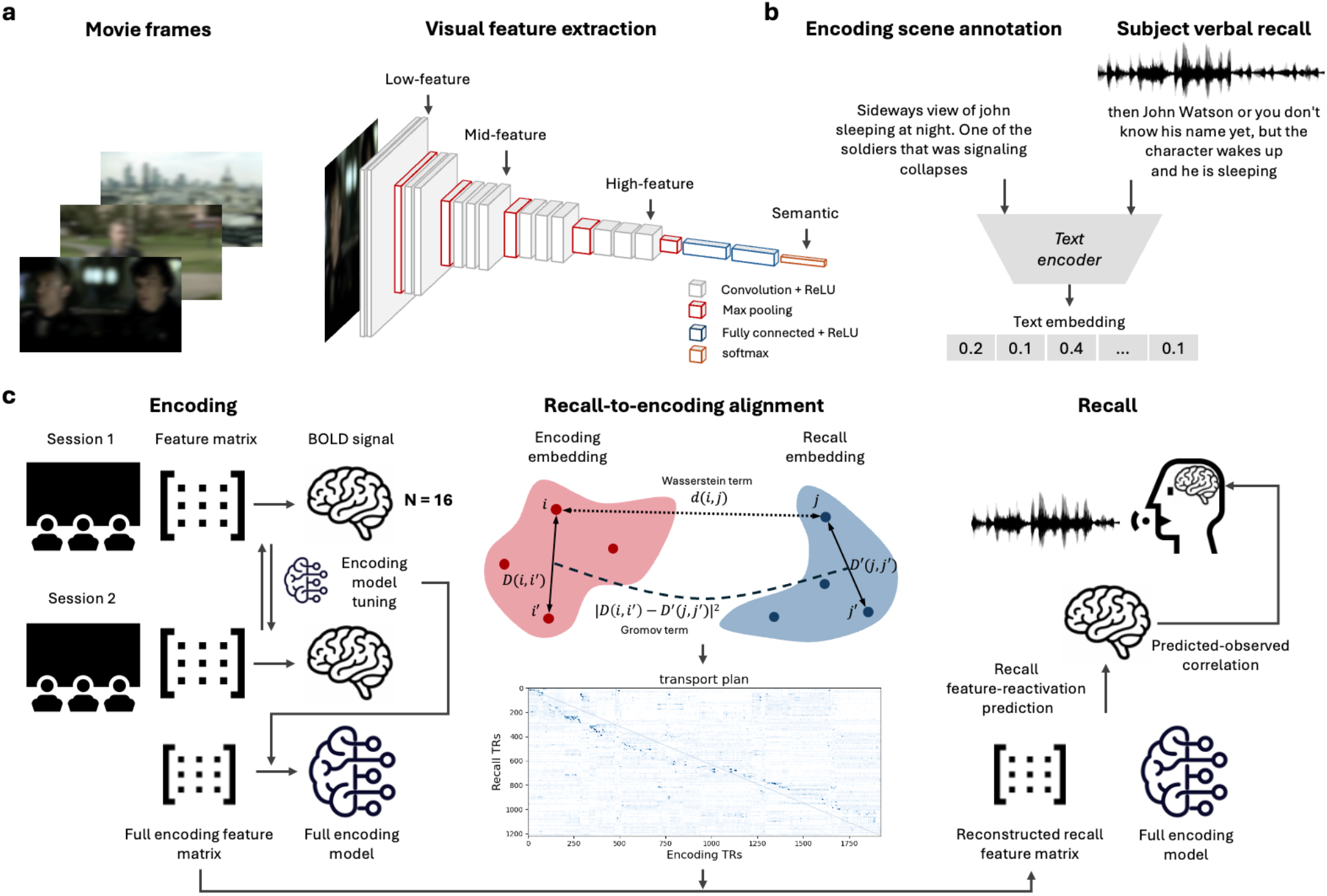
Optimal-transport alignment of movie encoding and free recall enables feature-resolved analysis of memory reinstatement. **(a)** Visual features were extracted from movie frames using a convolutional neural network (VGG-16), yielding a hierarchy of representations from low-level (early convolutional layers) through mid- and high-level visual features (layers 2, 7, and 13), plus a semantic feature from the final softmax output layer. Each feature level was reduced to its top 100 principal components to match input dimensionality across levels (see Methods). **(b)** Encoding scene annotations and subjects’ verbal recall transcripts were embedded into a common space using a text encoder of OpenAI’s *text-3-embedding-large* model, producing time-resolved text embeddings for both the viewed and recalled narrative. Further details on constructing per-TR text are given in Methods. **(c)** Analysis pipeline. For each parcel and VGG-16 feature level, cross-session prediction was used to select the number of latent components, and a final encoding model was fitted to both viewing sessions to predict recall activity. To align recall to encoding without prescribing event-to-event matches, recall and encoding embeddings were aligned by entropic optimal transport (middle panel): the transport plan maps each recall timepoint to encoding timepoints, providing a continuous, event-label-free viewing-to-recall correspondence derived from annotated text (the hyperparameters were selected by grid search under leave-one-subject-out cross-validation). The transport plan was used to reconstruct a recall feature matrix, which the encoding model then used to predict recall activity. Prediction accuracy was assessed as a rank-based accuracy derived from the correlation between predicted and observed recall responses.

We then asked whether recall preserved specificity of temporal information within events by testing whether predicted neural activation could distinguish different moments within the same manually segmented event. Next, we examined how prediction accuracy varied across the cortical hierarchy, using the principal gradient from sensory to association cortex^22^. A mixed-effects model tested the effects of state (encoding versus recall), feature level, and cortical gradient position. Finally, we quantified changes in feature-level differentiation between encoding and recall and used simulations to assess whether the observed convergence could be explained by temporal smoothing.

### The unsupervised alignment identifies recalled scenes and their encoding sources

To evaluate whether the transport plan captured meaningful correspondences between movie viewing and recall, we validated it against independent manual scene annotations that were not used to estimate the alignment. We conducted two complementary tests: whether the alignment distinguished scenes that participants recalled from those they omitted, and whether individual recalled segments were correctly aligned with their corresponding scenes in the movie.

First, we asked whether a given movie scene received greater alignment weight from participants who recalled it than from participants who omitted it. For each participant, we averaged the alignment weights across all recall TRs and across the encoding TRs belonging to each movie scene, yielding one alignment score per participant-scene pair. We then compared these scores, separately for each scene, between participants who recalled it and those who did not. This comparison holds scene identity constant, allowing us to assess whether alignment weights reflect individual recall rather than properties of the movie scenes themselves.

Of the 48 scenes considered, 41 were recalled by at least one participant. For 37 of these 41 scenes, alignment weights were higher among participants who recalled the scene than among those who omitted it (median difference = 1.77 × 10^−4^; one-sided Wilcoxon p < .001; Fig. 2a-b). The pattern remained when restricting the analysis to scenes recalled by at least three participants (25 of 26 scenes; median difference = 1.69 × 10^−4^; one-sided Wilcoxon p < .001). Thus, the alignment preferentially assigned weight to movie scenes that participants actually recalled, rather than distributing weights independently of the recall content.

**Figure 2.**
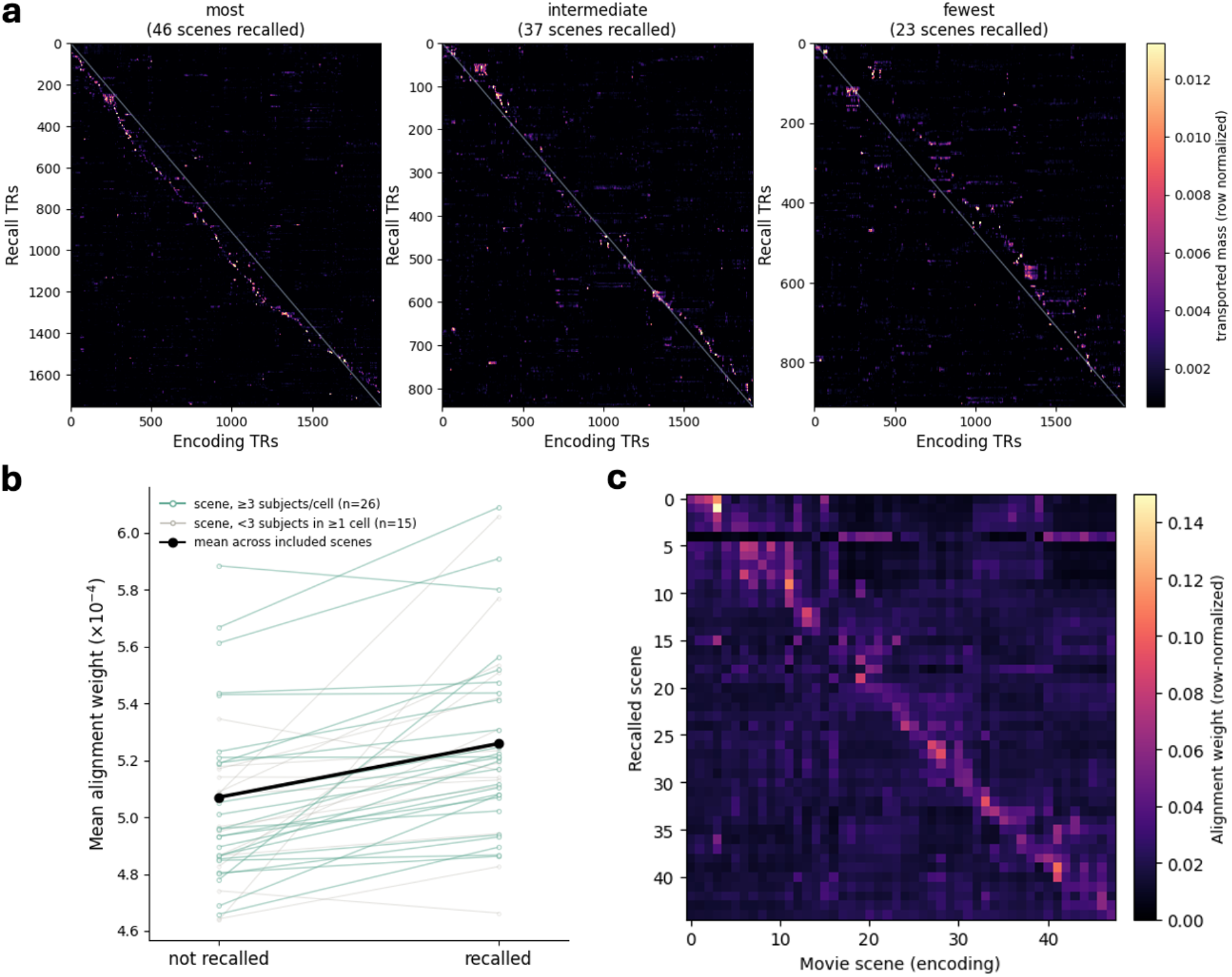
Behavioral validation of recall-encoding alignment. (**a)** Example transport plans for three participants who recalled the most (46 scenes), immediate (37 scenes), and the fewest (23 scenes) of the 48 movie scenes. Each plan maps recall TRs (rows) onto encoding TRs (columns); brighter cells indicate greater transported mass. Mass concentrates along the diagonal (grey line, 1:1 reference), falling below it where recall is temporally compressed relative to encoding. **(b)** Transported mass concentrated on recalled scenes. For each scene recalled by at least three and not recalled by at least three participants (n = 26), mean transported mass per TR was higher when the scene was recalled than when it was not (25/26 scenes positive; one-sided Wilcoxon p < .001). Green lines, included scenes; grey lines, scenes with fewer than three participants in either cell; black line, mean across included scenes. Because recalled-scene labels derive solely from the manual segmentation and never from the transport plan, all analyses reflect convergence between the unsupervised alignment and independent hand-labeling. **(c)** Scene-correspondence confusion matrix, summed across participants and row-normalized: for each recalled scene (rows), the distribution of transported mass across all movie scenes (columns). Movie scenes that were never recalled were omitted from the rows of the confusion matrix for visualization, but all 48 movie scenes were retained as candidate encoding sources when calculating identification accuracy. Mass concentrates along the diagonal (dashed line, correct correspondence). The unsupervised alignment recovered the correct movie scene well above chance (normalized rank accuracy 74%, chance 50%; median rank of the correct scene 7 of 48, chance is 24 of 48); exact-scene matching reached 16% (chance 2%) but errors fell on temporally adjacent scenes (peak mass within one scene of correct in 42% of cases), consistent with the loose temporal ordering of free recall.

Second, we tested whether the alignment could identify the specific movie scene corresponding to each recalled segment. For each manually segmented portion of a participant’s recall, we calculated its alignment weight with each of the 48 candidate movie scenes and ranked these candidates accordingly (Fig. 2c). The correct source scene achieved a mean normalized rank accuracy of 74% (chance = 50%), with a median rank of 7 among the 48 candidates. The correct scene ranked first in 16% of cases (chance = 2.1%) and among the top three candidates in 37% of cases (chance = 6.3%). In 42% of cases, the top-ranked scene was either the correct scene or an immediately adjacent scene (chance = 6.2%).

Together, these analyses demonstrate that the unsupervised alignment captures two complementary aspects of recall: which movie scenes participants remembered and where individual recalled segments originated in the movie. The alignment therefore provides a meaningful correspondence between continuous viewing and recall without requiring predefined scene boundaries.

### The reconstructed encoding features predict neural activity during recall

Having shown that alignment recovered meaningful behavioral correspondence between encoding and recall states, we next assessed whether this alignment captures information expressed in neural activity during recall. We first trained VGG-16-based feature specific encoding models for each participant, using the movie features to predict neural activation during viewing. Then we used each participant’s transport plan to reconstruct recall trajectories for each feature level and applied the corresponding trained encoding models to generate a predicted multivoxel activity pattern for each recall TR. Transport hyperparameters were selected using leave-one-participant-out cross-validation, so each participant’s recall prediction was obtained using parameters selected from the remaining participants (Supplementary Table S1). Prediction was evaluated by ranking the corresponding observed multivoxel pattern across all TRs within each state respectively, with 50% indicating chance.

Encoding models predicted movie-evoked activity above chance across all four feature levels, with widespread effects surviving FDR correction (Fig. 3a). Among all 300 parcels across feature levels, 90^th^ percentile rank accuracy ranged from 52.3% to 55.0%. Prediction for recall was weaker but remained reliably above chance across feature levels, with 90^th^ percentile rank accuracy ranging from 51.1% to 51.6%. Therefore, the reconstructed recall trajectories contained feature information the brain expresses during retrieval.

**Figure 3.**
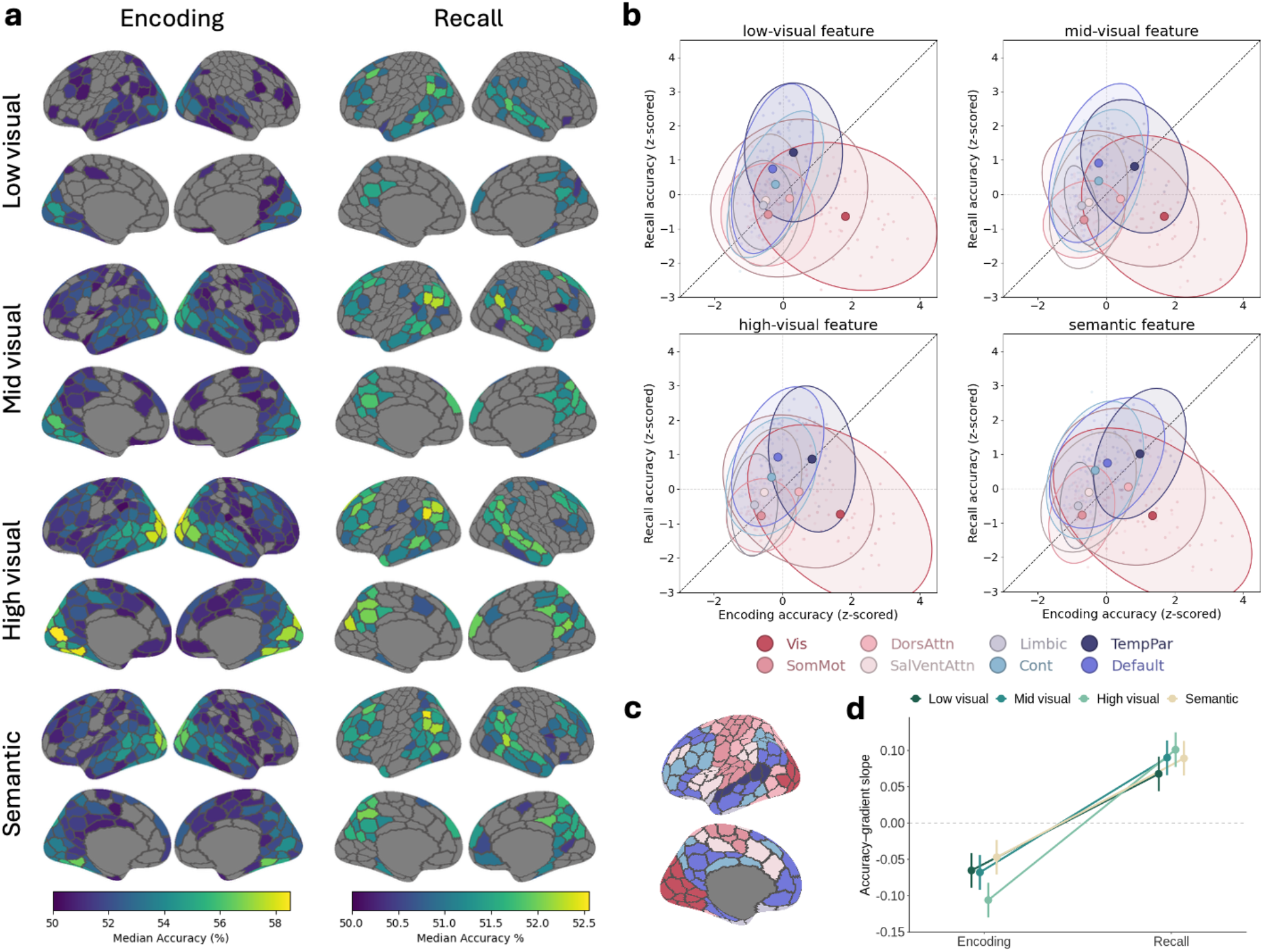
Feature-resolved reinstatement across encoding and recall. **(a)** Whole-brain maps of prediction rank-based accuracy (%) for each feature level (rows: low-, mid-, high-level visual, and semantic) at encoding (left) and recall (right). All parcels displayed on the map are at FDR-corrected q < .05 for each feature family. **(b)** Relative encoding versus recall accuracy across parcels by feature level. Each point is a parcel, colored by network. Parcels were segmented by the Yeo 17-network parcellation and grouped into eight networks (the Yeo-7 coarse networks plus temporoparietal). Vis = Visual; DorsAttn = Dorsal attention; TempPar: temporoparietal; SoMot: somatomotor; SalVentAttn: Salience/ventral attention; Cont: Control. Parcel accuracies are averaged across subjects and z-scored across parcels within each state, so the axes show relative standing rather than absolute accuracy. Filled markers are network centroids, with ellipses covering 95% of each network’s parcels. Dashed diagonal line marks equal relative standing. **(c)** The cortical distribution of the coarse network profiles defined in (b). **(d)** Slope of prediction accuracy along the principal cortical gradient (sensory to association)^22^, estimated from the mixed-effects model; error bars are 95% CIs. Positive values indicate higher accuracy toward association cortex, negative values indicate higher accuracy toward sensory cortex. Slopes are negative and feature-dependent at encoding, and shift positive and converge at recall.

### Recall reinstatement shifts from sensory to association cortex across feature levels

Having established that transported feature trajectories predicted recall activity, we asked whether their cortical distribution resembled those observed during encoding or was systematically reorganized during retrieval. For visualization, parcel-level encoding and recall accuracies were grouped based on their corresponding functional networks^23^. The distribution of parcels within each network was summarized using a 95% data ellipse.

Across feature levels, the encoding-to-recall relationship followed a consistent network organization (Fig. 3b-c). Visual parcels ranked relatively higher at encoding than at recall, whereas association parcels, particularly those in the default network, showed the reverse. This gross organization was apparent across all feature levels, where visual cortex was consistently encoding-dominant and the default network consistently recall-dominant regardless of feature levels.

To formally test this, we grouped the parcels according to their relative position in the cortical principal gradient^22^, in which lower values lean towards sensory cortex and higher values lean towards association cortex. We then quantified this redistribution by modeling rank accuracy as a function of state, feature level, and each parcel’s position along the gradient, including all interactions, crossed random effects for subject and parcel, and a by-subject random slope for state. Prediction differed overall between encoding and recall, F(1, 15) = 96.7, p < .001. Notably, its relationship with cortical position reversed between states, as indicated by a state × gradient interaction, F(1, 38032) = 1453.2, p < .001; β = 0.158, 95% bootstrap CI = [0.12, 0.19]. At encoding, prediction was stronger toward the sensory end of the gradient (β = -0.07, 95% CI = [-0.09, -0.05]), whereas at recall, prediction increased toward association cortex (β = 0.09, 95% CI = [0.07, 0.11]; Fig. 3d). Thus, both the network-level visualization and the gradient analysis showed a broad redistribution of stimulus-related prediction from sensory cortex during encoding toward association cortex during recall.

### Recall reinstatement carries temporal structure within events in association cortex

Event-level alignment in prior section established which narrative event is being recalled, but it does not determine whether recall activation also distinguishes different moments within that event. To test that, for each predicted recall TR, we restricted the candidate observed patterns to TRs from the same manually segmented recall scene. Because event identity was constant across candidates, above-chance rank accuracy could only arise from within-event temporal structure.

Within-event accuracy exceeded chance, indicating that the recall activity contained information about relative temporal position within manually segmented events. Significant parcels were concentrated in association cortex (Fig. 4a), while early sensory parcels were at chance despite being where encoding prediction was the strongest. To test whether this cortical redistribution was reliable across participants, we summarized each participant’s median within-event accuracy in sensory and association cortex, and tested their paired difference across participants. Median accuracy was 50.69% across association cortex versus 50.05% across sensory parcels with a reliable difference (median difference = 0.59%, 13/16 subjects, one-sided Wilcoxon p < .001; Fig. 4d).

**Figure 4.**
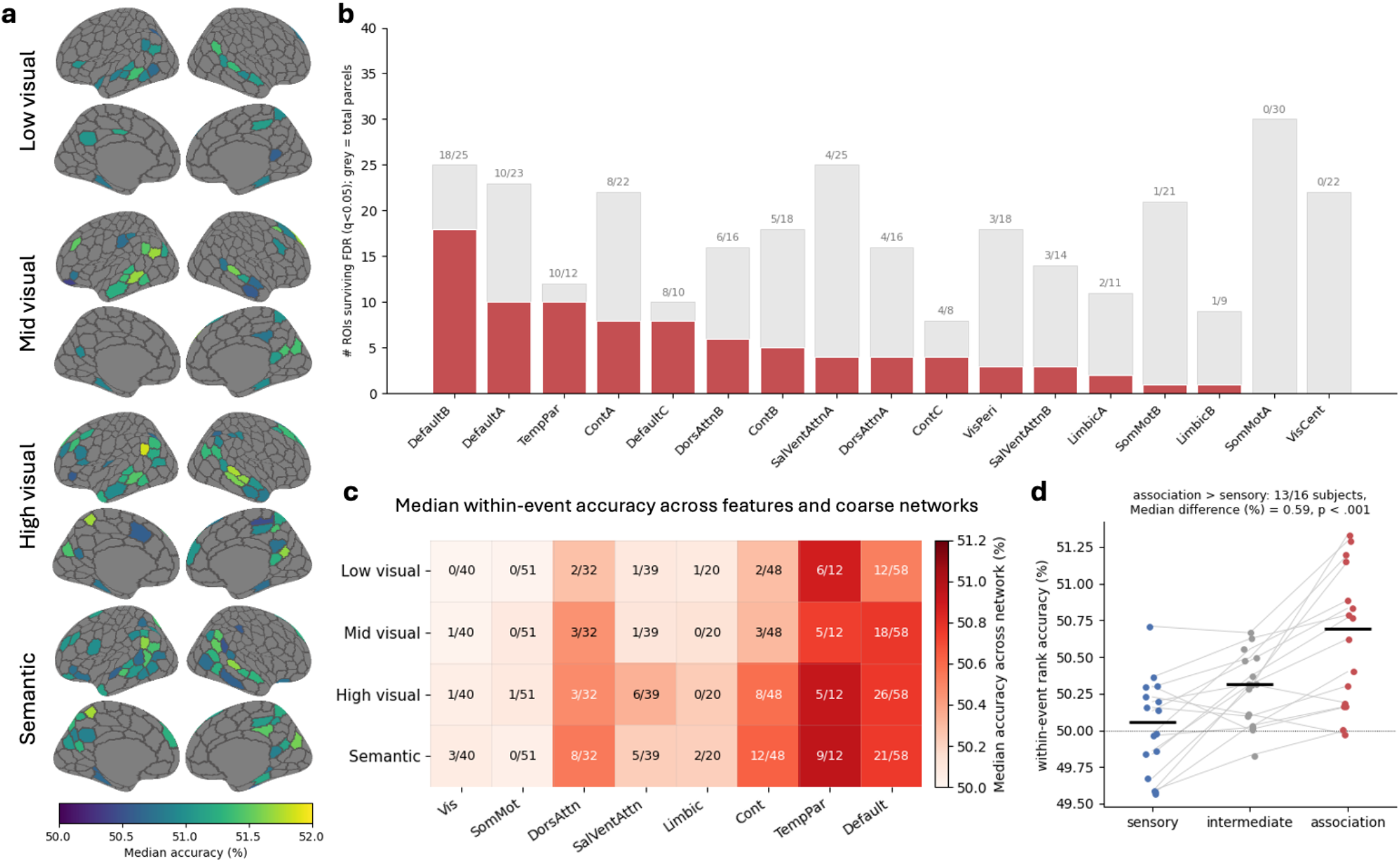
Within-event reinstatement is localized to association cortex. **(a)** Median within-event rank accuracy across subjects for each feature level (low visual, mid visual, high visual, semantic), projected onto the Yeo-17 network Schaefer 300-parcel atlas. Only parcels surviving FDR correction (q < .05) are shown. **(b)** Number of significant parcels within each Schaefer 17-network label (red), relative to the total number of parcels in that network (gray). Fractions are printed at each bar. Networks are ordered by the number of significant parcels. **(c)** Network-level within-event rank accuracy for each feature level across coarse cortical networks. For each subject, rank accuracy was summarized as the median across all parcels within each system, without thresholding by parcel wise significance, and then summarized across subjects. The numerator indicates how many significant parcels, and the denominator indicates the total number of parcels in this network. Cell color indicates group median within-event rank accuracy, with 50% corresponding to chance. The networks shown in the figure are a coarser-level of the 17-network labels (Vis, SomMot, DorsAttn, SalVentAttn, Limbic, Cont, TempPar, and the Default). **(d)** Subject-wise medians by cortical zone. Each point is one subject’s median rank accuracy across all parcels in a zone, averaged over the four feature levels. Gray lines connect the same subject across zones, black bars mark the group median, and the dotted line marks chance (50%). Medians are computed over all parcels in each zone, including those not surviving correction in (a). Networks were grouped as sensory (VisCent, VisPeri, SomMotA/B), association (DefaultA-C, TempPar, ContA-C), and intermediate (DorsAttnA/B, SalVentAttnA/B, LimbicA/B). Inference was computed in two stages. Within each subject we took the difference Medianassociation – Mediansensory, then tested those 16 difference scores against zero with a one-sided Wilcoxon signed-rank test (W = 129, p < .001; 13/16 subjects positive). The intermediate zone is shown for context and was not included in the contrast.

Intermediate networks fell between these extremes, consistent with a graded increase along the cortical hierarchy towards association cortex (Fig. 4b-c). Central visual and somatomotor A parcels were at chance. Only peripheral visual network showed a weak effect that remained below the association cortex’s level. These effects concentrated in regions that integrate information over longer temporal windows during narrative perception^8,9^.

A complementary full-window analysis was done to compare TR-resolved predictions with an event-averaged predictor (Supplementary Fig. S3). TR-resolved alignment produced significant increases in predictive performance over event-level prediction in association cortex, and these parcel-wise increments closely resembled within-event accuracy (Pearson’s r = .74). Thus, fine-grained alignment contributed information beyond coarse event identity, with both analyses converging on a preferential expression of this temporal structure in association cortex.

### Feature representations converge during recall along the principal cortical gradient

Here, feature separation refers to differences in predictive performance among feature levels, and convergence denotes a reduction in those differences. We tested whether separation decreased from viewing to recall using the state × feature interaction in the mixed-effects model described above, and whether that decrease varied along the cortical gradient using the three-way interaction. Full model statistics are reported in Supplementary Table S2. All accuracy points below are percentages.

The feature levels showed markedly less differentiation in prediction accuracy during recall than during encoding, a narrowing of between-level differences that we referred to as feature convergence (Fig. 5a, c). This pattern was reflected in a significant state × feature interaction, F(3, 38032) = 291.9, p < .001. At encoding, all six pairwise differences between feature levels were significant, ranging from 0.33 to 1.51 percentage points (95% CIs excluding zero). At recall, these differences shrank to 0.01-0.18 percentage points, and two comparisons – low versus mid-level and high versus semantic – were no longer significant.

**Figure 5.**
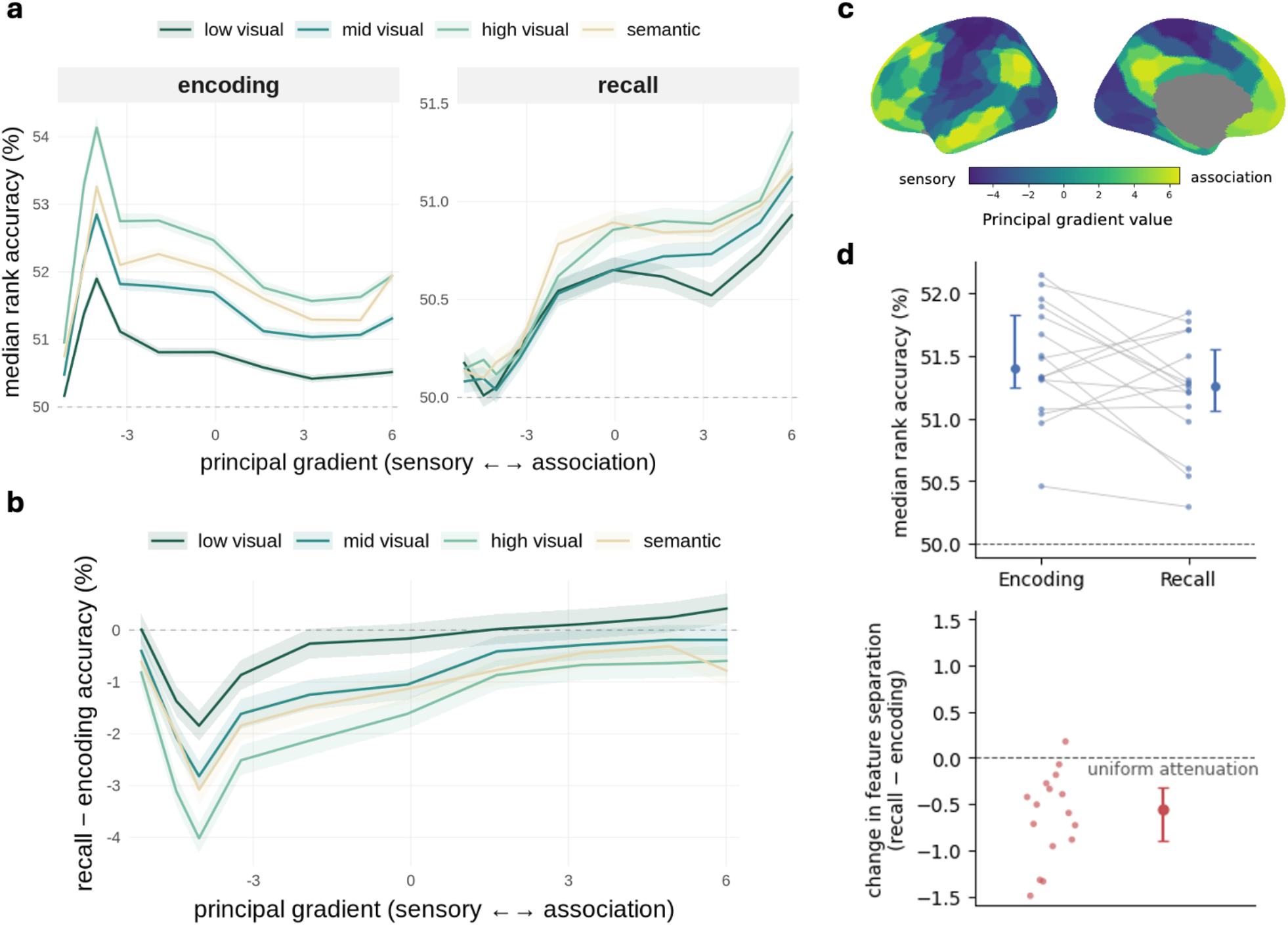
Feature separation is distinct at encoding and converges at recall along the principal gradient. (**a)** Median rank accuracy (in percentage points; 50 = chance) as a function of position along the principal gradient (binned into deciles), shown separately for encoding (left) and recall (right) and for each feature level (low-, mid-, high-level visual, semantic). Shaded bands denote ±1 SEM across parcels. At encoding, the four feature levels are clearly separated, with higher-level features predicting most accurately and accuracy declining from sensory toward association cortex. At recall, the feature levels converge and the relationship with the gradient reverses, with accuracy now rising toward association cortex. **(b)** Model-estimated recall-minus-encoding differences in median rank accuracy for each feature level across decile bins of the principal cortical gradient. For visualization of this change, the decile bins of the gradient was included as a categorical factor in an auxiliary mixed-effects model. This allows the encoding-to-recall difference to vary freely across the gradient without imposing a functional form. Points are plotted at the mean gradient value of parcels within each bin; lines connect bin-specific estimates, and shaded bands denote 95% confidence intervals. Statistical inference for the gradient interaction was based on the continuous-gradient mixed-effects model reported in the text. **(c)** The principal cortical gradient projected onto the cortical surface and parcellated for each parcel, running from sensory (low values) to association cortex (high values). **(d)** Median rank accuracy across parcels within the fixed union mask defined in recall (parcels showing evidence for at least one feature level at uncorrected p < .001; 78/300 parcels), averaged across all feature levels (top panel). The bottom panel shows the normalized feature separation between semantic and low-level features, calculated by their relative above-chance accuracies divided by their mean. Thin lines connect individual participants, with points and error bars showing the group median and interquartile range. The dashed line marks chance. Percentage is the median encoding-to-recall reduction, computed relative to chance for accuracy and relative to the encoding value for normalized feature separation. Because normalized separation is invariant to uniform scaling of the two feature signals, proportional weakening of recall prediction would leave this measure unchanged. Therefore, the observed reduction indicates convergence beyond overall attenuation.

To assess the convergence directly, we contrasted the semantic and low-level representations, which formed the endpoints of the sampled feature hierarchy. Their difference was 1.05 accuracy points at encoding, with 95% CI = [0.98, 1.11], but narrowed to 0.17 points at recall, 95% CI = [0.11, 0.23]. This reduction was reliable across subjects (β = -0.87, 95% bootstrap CI = [-1.07, - 0.70]).

This convergence was not a simple by-product of weaker recall prediction. Expressing separation relative to the mean above-chance prediction strength of the two feature levels yielded a measure invariant to uniform scaling of accuracy. Under uniform attenuation, normalized separation in recall should be the same as in encoding. Yet, normalized separation decreased from 0.788 at encoding to 0.200 at recall (Wilcoxon signed-rank, one-sided p < .001; Fig. 5d), indicating a loss of feature separation beyond the overall reduction in predictive performance. Sensitivity analyses confirmed that this effect was not driven by near-chance recall endpoint signals, and the reduction was in the same direction across all six feature-level comparisons (Supplementary Tables S3-S4).

An alternative is that the observed feature convergence is induced by the optimal-transport alignment. Because each recall time point is reconstructed from a weighted combination of encoding time points, the alignment may average together features from different moments in the movie, reducing their temporal specificity. To assess whether this smoothing could explain the convergence effect, we first examined local temporal mixing using Gaussian smoothing to the predicted recall activity, systematically varying the width of the smoothing kernel (Supplementary Results S2). We identified the kernel width that reproduced the degree of feature convergence observed with the OT alignment, then applied the same amount of smoothing to encoding predictions to determine whether it produced a comparable reduction in feature separation. Results suggested encoding separation remained approximately four times that at recall (0.796 vs. 0.200; 14/16 participants, two-sided Wilcoxon p < .001; Supplementary Fig. S4a-d). We then examined non-local mixing by applying temporal weight distributions derived from the fitted OT plans to encoding features. Despite substantial contributions from temporally distant movie moments, normalized feature separation increased from 0.788 to 1.087, remaining greater than during recall (0.200; 15/16 participants, two-sided p < .001; Supplementary Fig. S4e-f). Thus, temporal imprecision caused by our alignment was insufficient to reproduce the feature convergence observed during retrieval.

The feature convergence also varied along the cortical gradient, as indicated by a state × feature × gradient interaction, *F*(3, 38032) = 16.9, p < .001 (Fig. 5b). The encoding-to-recall change in gradient slope was larger for high-level visual features (0.21 accuracy points per unit gradient increase) than for other levels (0.13-0.16). This difference was reliable only between high- and low-level features, β = 0.08, 95% bootstrap CI = [0.05, 0.10], with the rest indistinguishable from one another. Thus, feature levels differed modestly in their encoding-to-recall change, while the graded separation present at encoding was largely reduced.

Together, recall involved two related but distinguishable changes: a broadly shared redistribution towards association cortex, and a substantial convergence of feature-level organization.

## Discussion

The present study asked whether memory reinstates feature-specific representations from the *Sherlock* television episode during continuous narrative recall. Prior naturalistic-memory work concerning neural reinstatement treated events as unitary patterns^10,11,12,13^, without considering the continuously-evolving information contained within events. We recovered the correspondence between movie viewing and free recall with a customized framework of optimal transport, which requires no manual event labels or predefined boundaries. We then used feature-specific encoding models trained on viewing data to test whether the resulting temporally resolved feature trajectories predicted recall activity. Preserving variation within manually defined events improved prediction primarily in default-mode and frontoparietal cortex, systems previously linked to event-level reinstatement in this dataset^10,13^, indicating that reinstatement retained temporal specificity of moments within events. Retaining this temporal resolution further allowed us to characterize how the constituent features of narrative experience are organized during recall. We found that the cortical distributions of feature levels changed systematically along the principal cortical gradient, with the largest differences for high-level visual features. Feature separation was also substantially reduced during recall, producing a convergence of predictive performance across feature levels. Together, these findings suggest that narrative recall is not simply a weakened replay of perception, but a reconstruction that preserves the unfolding structure of events while reorganizing their constituent information into a more integrated cortical representation.

### Within-event organization of naturalistic recall

Research on naturalistic perception has shown that people spontaneously divide continuous experience into meaningful events. Event segmentation theory (EST) proposes that continuous experience is segmented when changes in ongoing input trigger updates to an event model^24,25^, which in turn shapes the organization of memory as expressed at recall^26,27,28^. Accordingly, individual scenes from a TV show are reinstated during free recall in posterior midline and default-network regions^10,11,29^, and perception-derived event states recur during retrieval^12^.

Reinstatement has also been demonstrated for shorter movie segments linked to individual recall utterances^13^. However, neural similarity between corresponding viewing and recall segments does not necessarily establish whether recall activity distinguishes different moments within the same event. For example, recalling Sherlock examining a body might evoke a neural pattern similar to the corresponding movie segment because both reflect the broader crime-scene event, without distinguishing that moment from his arrival at the scene.

Our findings show that, by aligning continuous viewing and recall trajectories without predefined event segment correspondences, cortical regions previously implicated in event-level reinstatement carry information that distinguishes different moments within the same event.

Retaining within-event variation improved prediction accuracy over and above event-level models, particularly in default-mode and temporoparietal networks associated with contextual and situation-level representations^30,31^. Behavioral work likewise indicates that naturalistic recall preserves organization below the level of coarse event sequences. Prior work has shown that narrative recall preserves the broad sequence of events despite variability in the details remembered^32^. Similarly, recall of real-world experiences retains temporal structure, with stronger temporal organization associated with richer episodic detail^33^.

The concentration of this within-event information in higher-order association cortex may initially seem counterintuitive, as these regions were generally associated with relatively long temporal receptive windows^8,9^. However, the timescale over which a region integrates information is different from how dynamically its represented content can change. Indeed, higher-order cortical responses can depend on extended narrative context while flexibly updating or discarding prior contextual information^34^. Theories of lateral parietal cortex similarly characterize it as an online buffer that maintains temporally extended context while updating its contents as ongoing information changes^35^. Consistent with this distinction, dorsomedial posterior parietal neurons in macaques were found to be sensitive to time-varying events and sensory features while also accumulating information over timescales extending to tens of seconds^36^. Our findings further suggest that association cortex supports moment-resolved event coding, in which neural representations distinguish changing moments within an extended narrative event rather than reducing the event to a single, undifferentiated state. Thus, long temporal receptive windows need not imply coarse temporal specificity, suggesting that integration over extended contexts and differentiation of momentary contents can coexist during naturalistic recall.

### Feature organization converges during naturalistic recall

A central question in the reactivation literature concerns whether information represented at encoding reinstates at retrieval. Neural reactivation has been demonstrated for categories, items, events, perceptual attributes, and multiple feature levels across sensory and frontal cortex^1,2,3,37,38,39,40^. Yet the presence of the same information during perception and retrieval does not necessarily imply that it is represented in the same way^41,42^. Recall might simply weaken the feature-specific signals observed during perception, leaving their relative strengths and cortical distributions intact, or it might change how these signals are expressed across the cortex. We distinguished these possibilities using two properties of feature level organization. First, we measured the cortical profile of predictive performance for each feature level along the sensory-to-association gradient^22^. Second, we measured feature separation, defined as the differences in predictive performance among feature levels when evaluated at the same cortical position.

Our results were inconsistent with uniform attenuation of the feature-level organization present during viewing. Under such an account, above-chance predictive performance would be reduced by a similar proportion across feature levels, preserving their relative separation after accounting for the overall reduction in signal strength. Instead, normalized feature separation decreased substantially even though individual feature levels remained reliably predictable. The cortical profiles of these signals also changed. The magnitude of attenuation differed across feature levels, and the relationship between predictive performance and position along the sensory-to-association gradient differed between viewing and recall. Thus, recall did not simply weaken the feature-level signals present during viewing. It reduced the separation among feature levels while reorganizing how their predictive information was distributed across cortex.

Related findings from episodic retrieval suggest that internally generated representations can preserve stimulus information while reducing distinctions that characterize perception. Color and object information were found to be more separable during perception than at retrieval^5^, and the strong gradient in spatial precision across the visual hierarchy during perception was not reproduced during memory retrieval^7^. A similar reduction in perceptual differentiation has also been observed during visual imagination, where representations of imagined objects were more similar to one another than representations of perceived objects^43^. More broadly, this pattern fits transformation accounts in which remembered information remains detectable while changing systematically in where or how it is expressed relative to perception^4,6,44^.

Together, these findings suggest that recall reorganizes rather than simply attenuates the feature-level structure established during viewing. One possible contributor to this convergence is greater top-down selection during retrieval. Even during perception, goal relevance can flexibly alter feature representations, particularly in frontoparietal networks^43^. During recall, such selection may operate within contextual or schematic representations that determine which aspects of an event are currently relevant^46,47,48^. This may be especially relevant for narratives, in which later information changes the relevance of earlier moments. Consistent with this idea, narrative transitions can reactivate contextually relevant information from earlier events^47^. These shared top-down constraints, as a scaffold, could reduce feature separation while preserving temporally specific information. Our findings therefore extend prior evidence for perception-to-memory transformation by showing that, during recall of an extended narrative, feature-level reorganization coexists with preserved temporal specificity of varying contents within events, consistent with flexible expression of memory representations at different levels of specificity^18^.

### Limitations and future directions

Two limitations exist in this study. First, we derived the feature levels from successive network layers that are correlated with one another and do not correspond exactly to stages of human visual processing, particularly at higher levels^50^. However, because viewing and recall are evaluated in the same model-defined feature spaces, this limitation does not undermine our comparison of these same features between these two states. Second, the analyses were predictive rather than reconstructive. Future work could apply generative models directly to recall activity and compare it with reconstructed recall activity through this transport plan method, which may provide a more detailed assessment of content transformation account.

Moreover, as the transport plan is estimated separately for each participant, future work could also examine whether properties of the alignment relate to stable individual differences in retrieval style^51,52^. Relating these profiles to participant-specific transport properties could test whether such trait differences are expressed in the dynamics of naturalistic recall.

## Conclusion

Real-world experiences unfold continuously, while memory studies have typically examined isolated stimuli or treated extended narrative events as unitary representations. Here, we modeled the temporal evolution of a remembered narrative without imposing any predefined event boundaries, revealing that recall activity in the association cortex can distinguish changing contents within individual events, even as the feature-level organization that characterized those viewing moments was substantially reorganized. Thus, becoming less perception-like does not mean becoming less temporally specific. These findings suggest that narrative memory can preserve *how* an experience unfolded without preserving the perceptual organization of all its contents.

## Methods

### Dataset

The data used in this study was from an fMRI study by Chen et al.^10^, where 16 subjects watched the first 48 minutes of the first episode of *BBC’s Sherlock*, and then freely recalled them while being scanned. The movie watching was divided into two sessions (23 and 25 minutes) to reduce the length of each individual runs. Participants received specific instructions for the verbal recall task: 1) recall the movie in its original sequence whenever possible; 2) prioritize completeness and detail over strict temporal order.

The experiment was conducted in a 3T scanner using a T2*-weighted echo planar imaging (EPI) pulse sequence (TR = 1,500 ms, TE = 28 ms, flip angle 64°, whole-brain coverage with 27 slices of 4 mm thickness, in-plane resolution of 3 x 3 mm, with FOV of 192 x 192 mm), in interleaved acquisition.

### MRI data preprocessing

All MRI data were applied with same preprocessing steps through AFNI software^53^ using afni_proc.py. Voxel time series were despiked (*despike*) before slice-timing correction (*tshift*). The corrected functional images were then aligned to the anatomical scans (*align*) and subsequently normalized to MNI space (*tlrc*). Additionally, volume registration (*volreg*) was executed by aligning functional volumes to a reference volume identified as having the least outliers, followed by a spatial smoothing with a Gaussian kernel of 4 mm full-width at half-maximum (FWHM) (*blur*). Covariates considered as nuisance factors were regressed out from the functional data. Specifically, these included mean signals derived from two eroded masks representing ventricles and white matter, the global signal, and a 24-parameter motion regression model encompassing six basic motion parameters, their temporal derivatives, and respective squared terms^52^. To preserve the temporal integrity essential for this continuous movie dataset encoding model training, no volumes were censored.

Voxels were then grouped by Schaefer-Yeo-17 network 300-parcel functional brain atlas^19,23^. The atlas was downsampled to match the resolution of the MNI-registered functional data (3 ξ 3 ξ 3) with *3dresample*.

### Movie data preprocessing

Frames from movie data were processed by *ffmpeg* Python package with frequency of 2 Hz with a frame size of 224 x 224 for alignment with the expected input size of the vision model (see below) used to extract image features. To align with the sampling resolution of the fMRI data, each frame was passed through VGG-16^20^ (Simonyan & Zisserman, 2015), a convolutional neural network (CNN) that extracts increasingly abstract visual features across successive layers. VGG-16’s and related CNNs’ successive layers have demonstrated strong correspondence with brain activity in visual cortex^14,15^. To extract feature-level activations, we followed Bone et al.^3^, who sampled VGG-16 layers representing progressively higher levels of visual information. For each frame, we extracted activations from convolutional layers 2, 7, and 13 and the final softmax output layer as approximate representations of low-, intermediate-, high-level visual, and semantic features, respectively.

For each input image, activation maps were extracted from the selected convolutional layers. Specifically, each channel-specific activation map was downsampled to a 3 x 3 spatial grid using area-based interpolation to account for the limited retinotopic precision associated with participants’ eye movements^3^. The resulting maps were flattened and concatenated across channels, producing feature vectors of 576 dimensions for the low-level layer (3 x 3 x 64), 2304 dimensions for middle-level, 4608 for high-level. The semantic representation consisted of the 1,000-dimensional output over ImageNet categories and therefore was not spatially downsampled. These per−layer feature vectors were averaged per TR, followed by a z-standardization on the training vectors, and applied to both training and test matrices to get zero-mean, unit-variance standardization. We applied dimensionality reduction via principalcomponent analysis (PCA) separately to each layer’s feature matrix for each movie part and projected on to the other movie part, retaining the top 100 components per layer. PCA compression to a fixed number of components adjusted for differences in raw feature dimensionality across layers and ensured that each encoding model had matched input size for comparison.

### Visual Encoding Model Building and Testing

To train feature-specific visual encoding models, we used beta estimates of fMRI activity at the per-TR level and the corresponding feature activations derived from the CNN model at corresponding time-points. To account for the hemodynamic response function (HRF) delay while allowing for region-specific delay variability (typically 4-6 seconds), we computed feature matrix by shifting the model in time by 3 TRs (4.5 s). For each node, we fit a partial least squares (PLS) regression using *scikit-learn* package in Python to predict neural activity from feature activations. We let *β* _i,t_ denotes the beta estimate for node *i* at time *t*:

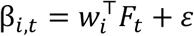

The ***w****_i_*^T^ stands for the learned PLS regression beta weight vector for node *i*. ***F****t* is the feature vector at timepoint t, *χ* stands for residual noise.

To evaluate encoding model performance without circularity, we used a bidirectional cross-session procedure. First, the model was trained on encoding session 1, with three-fold cross-validation conducted within that session to select the number of latent components for each parcel. The optimized model was then used to predict the multi-voxel activity pattern in encoding session two. This procedure was repeated in the opposite direction. Prediction accuracies from the two train-test directions were averaged to provide an estimate of overall model performance.

The model performance was quantified as correlation-based rank accuracy. For each parcel, we computed a correlation matrix between the predicted and true multi-voxel patterns across TRs in the held-out session. For each predicted TR, all observed TRs were ranked according to their correlation with the predicted pattern, and the rank of the temporally corresponding observed TR was recorded. These ranks were normalized and averaged across TRs, then expressed as percentages, with 100% indicating perfect identification and 50% as chance level.

Statistical significance was assessed using a null distribution generated from 1,000 block permutations of the predicted encoding neural time series. Each block consists of 20 TRs, which preserves temporal autocorrelation while disrupting its correspondence with the fixed predicted patterns. For each participant, parcel, and feature, we calculated the median rank accuracy across permutations. We then subtracted this empirical null’s median from the observed rank accuracy.

Rank accuracy was estimated by two-fold cross-validation across the two movie parts. For each participant, parcel, and feature, null median-centered accuracies were averaged across the two train-test directions to yield a single estimate, and which was then tested against zero across participants using a one-sided one-sample Wilcoxon signed-rank test. Benjamini-Hochberg false-discovery rate correction (FDR) was then applied across p values across parcels within each feature level.

### Text embedding extraction

To build an alignment space between movie encoding and recall, we used the movie scene annotation and individual subjects’ word-level recall transcripts with timestamps^10,21^.

Movie annotation was documented as segments with varying time duration, nested under manually defined coarse scene segments^10^. Overlapping or duplicate segments were resolved before analysis. Specifically, exact duplicates were merged, and the start time of each remaining overlapping segment was shifted to the next first non-overlapping time point. Each scene was characterized by its annotated event description together with contextual information (indoor/outdoor, named characters, and location). For each TR, we used *GPT-4o* (temperature = 0) to merge the scene’s context and description into a single prompt (Supplementary Material S1). Each sentence was then embedded with OpenAI’s *text-embedding-3-large* (3072 dimensions) to obtain its semantic embedding.

To provide sufficient sentence-level semantic contents per recall TR, we required each TR to contain at least 15 words, a threshold chosen to approximate the scale of contextual information previously shown to support neural language representations^55,56^. To do that, we augmented the TRs with fewer words with subsequent words, producing a semantically smoothed raw texts per TR. The resulting text was then processed by the same text encoder to extract a semantic embedding.

### Recall to encoding mapping

Because free recall involves selecting, integrating, compressing an extended event^18^, and memory retrieval unfolds over continuous events rather than isolated moments, mapping recall to encoding requires a soft alignment across time rather than a rigid point-to-point match. To achieve this, we modeled each recall timepoint (TR) as a soft probability distribution over encoding TRs using optimal transport (OT) through Python package *POT*^57^. Specifically, we used fused Gromov-Wasserstein distance (FGW)^17^ to balance two complementary alignment goals governed by *α*: matching features directly across encoding and recall (the Wasserstein term: *what* content is retrieved), and preserving relationships among timepoints within each state’s representational geometry (the Gromov term: *how* that content is organized; Fig. 1c).

These relationships were defined using cosine distances between alignment embeddings.

For recall and encoding feature distributions with samples *x_i_* and *y_j_*, where *i and k* index recall TRs and *j* and *l* index encoding TR, intra-domain structure matrices *C^x^*, *C^y^*, and a cross-domain feature cost *M_i,j_*, the entropy-regularized OT-FGW transport plan is thus:

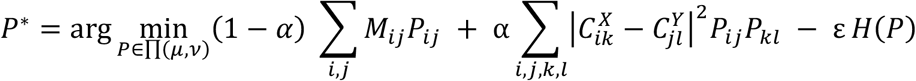

Where ∏(*μ*, *ν*) is the set of couplings with the prescribed recall and encoding marginals, *H*(*P*) = − ∑*_ij_ P_i,j_* log *P_i_*_,*j*_ is the Shannon entropy, *ε* controls the regularization, and *α* ∈ [0, 1] trades off the Wasserstein feature term against the Gromov structure term.

Because the effective influence of the feature, structural, and entropy terms depends on their relative numerical scales, we rescaled the cost matrices separately for each participant before fitting FGW. The within-domain cosine-distance matrices *C^X^*and *C^Y^* were each divided by the median of their off-diagonal entries, whereas the cross-domain feature-cost matrix *M* was divided by the median of all its entries. These rescaling placed the feature and structural costs on comparable scales before they were weighted by *α*. Finally, we z-scored each encoding-TR column across recall time to express alignment relative to that encoding TR’s typical coupling strength, then applied a softmax across encoding TRs within each recall TR to obtain a normalized recall-to-encoding alignment distribution (inverse temperature = 1).

Before analyzing the recall data, we fit a final encoding model using data from both encoding sessions. Unlike the cross-session evaluation described above, in which model dimensionality was selected by within-session cross-validation, here we used cross-session prediction solely to select the number of latent components. In each direction, candidate models were trained on one encoding session, and the number of components that maximized prediction accuracy in the other session was selected for each parcel and feature level. We then averaged the two selected component counts and used this value to fit the final encoding model on the combined data from both sessions.

To predict recall data, for each feature level, we reconstructed recall-time features by multiplying the encoding feature matrix with the OT transport plan. The reconstructed features were then passed through each participant’s encoding model to generate predicted neural activity at every recall TR. Prediction performance was evaluated as described above: at each TR, the predicted multivoxel pattern was correlated with the observed pattern, converted to a rank among candidate time points, and expressed as normalized rank accuracy.

Hyperparameters of the transport plan were selected using leave-one-subject-out cross-validation through grid search (Supplementary Table S1; Fig. S1). For each grid, prediction accuracy was averaged across parcels within each participant and then across the training participants. The combination yielding the highest mean accuracy was selected and applied to the held-out participant. Recall features and neural predictions for that participant were then recomputed using the selected hyperparameters.

### Behavioral validation of the alignment

To assess whether the OT plan recovered meaningful correspondence between encoding and recall, we validated it against the manual scene-level segmentations of both the movie and each subject’s recall^10^, with a total of 48 encoding scenes (two unrelated cartoon scenes were removed). We performed two tests. The first asked whether the alignment distinguished movie scenes that an individual participant recalled from those they omitted. The second asked whether individual recall segments were aligned specifically to their corresponding source scenes.

For each subject *i* we quantified the alignment with movie scene *s* as

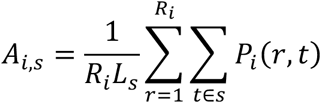

Where *P_i_*(*r*, *t*) is the normalized alignment probability from recall TR *r* to encoding TR *t*, *R_i_* is the number of recall TRs for participant *i*, and *L*_3_ is the number of encoding TRs in movie scene *s*. Because each recall TR was represented by a normalized distribution over encoding TRs, averaging across recall time makes alignment values comparable across participants with different recall durations.

Using the manual segmentation of each subject’s recall, we labeled each encoding scene as recalled or not-recalled for that subject. These labels derive only from the annotations, never from the transport plan, so agreement between mass and labels reflects convergence of two independent procedures.

To test whether alignment tracks individual differences in recall rather than overall scene salience, we restricted to scenes recalled by at least one participant. For each scene, we compared the mean alignment weight between subjects who recalled it and those who did not, and tested the scene-level difference with a one-sided Wilcoxon signed-rank test. Pairing by scene holds scene-level properties (length, position, salience) constant.

This analysis thus establishes whether recalled scenes receive greater alignment overall, while which has not yet answered whether individual recall segments are mapped to their specific source events, a matter of precision. To resolve that, for each recalled scene *k*, we calculated its alignment with every candidate movie scene *s*:

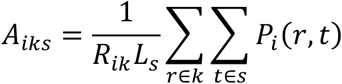

Where *R_ik_* is the number of recall TRs belong to event k. Thus, different from the preceding analysis, alignment was averaged only over the portion of recall corresponding to that particular recalled event. We then ranked all 48 movie scenes by their alignment score and recorded the rank of the manually corresponding source scene. We summarized the corresponding using the rank of the correct (same-identity) movie scene. We reported normalized rank-accuracy (chance = 0.5), the median rank of the correct scene, the exact-match (top-1) rate, and the proportion of recalled scenes whose peak mass fell within one scene of the correct scene. The candidate pool for ranking comprised all 48 movie scenes, including scenes recalled by no participant.

### Neural validation of the alignment

To test whether the OT alignment recovered correspondence expressed in recall neural activity, we asked whether movie features transported into recall time predicted the observed multivoxel activity pattern at each recall TR. Statistical significance was assessed in the same way as in the encoding data prediction, despite here we only tested on recall data.

The OT alignment was compared against a diagonal baseline, in which each recall TR was matched to the encoding TR at the same relative position, assuming recall linear chronological replay of encoding. The baseline used the same feature reconstruction and neural readout, so the two approaches differed only in the alignment step. OT yielded higher accuracy than the diagonal baseline at all feature levels (Supplementary Results S1).

To assess whether TR-resolved alignment predicts recall activity beyond what event-resolved alignment achieves, we repeated the identical pipeline with a single change to the predictor. For the coarse condition, each predicted recall pattern was replaced by the mean predicted pattern across all recall TRs assigned to the same annotated event. The coarse predictor is therefore constant within an event and cannot discriminate among that event’s TRs, while the transport plan, encoding models, and observed recall data are held fixed. Both predictors were scored identically, where every recall TR remained a separate query, ranked by pattern correlation against all observed recall TRs. The TR-resolved increment was defined as the difference in rank accuracy between the TR-resolved and event-averaged predictors. Statistical significance was assessed using a paired null distribution generated by applying the same block permutation to both conditions. Increments were tested across participants for each parcel and feature using one-sided Wilcoxon signed-rank tests, with FDR correction across parcels within each feature level (Supplementary Fig. S3).

### Within-event analysis

The comparison with an event-averaged predictor tests whether retaining temporal variation improves prediction across the full recall time series. It does not isolate within-event information, because the improvement can reflect better discrimination both between and within events. To test within-event structure directly, we restricted each query’s candidate patterns to TRs from the same manually labeled scene. Scene identity was therefore held constant, allowing the ranking to assess whether the predicted pattern distinguished moments within that scene.

Within-event rank accuracy was computed by ranking each query TR against the observed patterns of all TRs in its own scene and averaging the normalized ranks across all scored TRs. The null distribution was generated by permuting query rows within each scene over 1,000 permutations, which removes within-event position while keeping scene membership intact. Null-centering, the group one-sided Wilcoxon test, and FDR correction followed the procedure above.

### Cortical gradient parcellation

To situate parcel-wise effects along the cortex’s principal functional axis, we used the first connectivity gradient^22^, which spans a continuum from primary sensory/motor regions to association cortex. The dense gradient map was obtained in fsLR space at 32k resolution via the *neuromaps* toolbox^58^. To express the gradient at the resolution of our functional analyses, we parcellated it using the Schaefer-Yeo 17-network 300-parcel atlas in fsLR space, assigning each parcel the mean gradient value across its constituent vertices via the function *Parcellator.* This yielded a single principal-gradient score for each of the 300 parcels, aligned to the parcel ordering used throughout the neural analyses.

### Quantifying the transformation of feature-level representations from encoding to recall

To characterize how feature representation changed from encoding to recall, we fit a linear mixed-effects model to the per-parcel rank accuracies (chance = 50%):

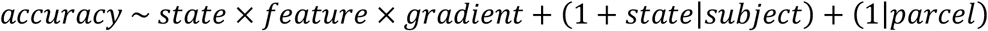

Feature level was coded with sum-to-zero contrasts and state with encoding = 0 and recall = 1. Gradient was mean-centered across parcels. In this setup, the state × feature interaction tests whether the feature-level profile differs between encoding and recall at the mean gradient position. The state × feature × gradient interaction tests whether this encoding-recall difference varies along the cortical gradient.

Models were fit by REML in R with *lme4/lmerTest* package, with fixed effects tested using Type III F tests and Satterthwaite-approximated denominator degrees of freedom. Because the state varies within subject and carries a by-subject random slope, the denominator df reflects the number of subjects. The other terms could not be given by-subject slopes without convergence failure at this sample size and were therefore tested against residual rather than subject-level variance, making the p value anticonservative. We refit the primary model on 500 subject-level cluster bootstrap resamples and report 95% bootstrap confidence intervals (CI) for the reported fixed effects. To assess how each feature behaves for each state and in cortical gradient, pairwise comparisons were estimated from the primary fit using estimated marginal means (*emmeans*) with Wald 95% CIs.

### Feature convergence quantification

The mixed effect models above could not assess whether the convergence would be due to simply a weaker recall signal, to quantify this convergence, we contrasted the low-level and semantic features, two endpoints of the feature hierarchy. To allow for a direct comparison between encoding and recall, we restricted the analyses to a union mask defined as the parcels showing above-chance significance in any feature level during recall (uncorrected p < .001; 78/300 parcels). We then applied this same mask to encoding and recall to calculate the feature separation.

To obtain feature separation, for each participant and state, we calculated the median rank accuracy across parcels within this mask for low-level and semantic features and subtracted chance (0.5). Normalized feature separation was defined as

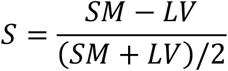

where LV and SM denote the above-chance low-level visual and semantic accuracies, respectively. This measure is invariant to uniform multiplicative scaling of the two feature signals, so proportional attenuation of both signals alone leaves normalized separation unchanged. The same mask and normalized feature separation were also used for supplementary analyses for blurring controls (Supplementary Results S2). A one-sided Wilcoxon signed-rank test was used to assess whether there exists any significant feature convergence beyond the proportional attenuation.

## Data availability

The dataset analyzed in this study are available on OpenNeuro (id: ds001132).

## Code availability

The pipeline codes, figure plotting codes with derived data to replicate the key findings in this paper are available on GitHub: https://github.com/shikangpeng/sherlock-OT.

## Acknowledgement

Computations were performed on the Trillium supercomputer at the SciNet HPC Consortium. SciNet is funded by Innovation, Science and Economic Development Canada; the Digital Research Alliance of Canada; the Ontario Research Fund: Research Excellence; and the University of Toronto.

## Author contributions

Conceptualization: S.P., B.L., B.R.B.

Methodology: S.P., B.R.B.

Data curation, formal analysis, & visualization: S.P.

Supervision: B.L., B.R.B.

Writing – original draft: S.P.

Writing – review & editing: B.L., B.R.B.

