## Supplementary for "Naturalistic recall preserves temporal specificity despite cortical feature reorganization"

### Outline

| | | |
|---|---|---|
| **Table S1.** | Hyperparameter search space and cross-validated selections | pg. 2 |
| **Figure S1.** | Cross-validated sensitivity to entropy regularization | pg. 2 |
| **Results S1.** | OT outperforms diagonal mapping | pg. 3 |
| **Figure S2.** | OT alignment outperforms a diagonal baseline | pg. 3 |
| **Figure S3.** | Temporal granularity comparison for recall | pg. 4 |
| **Results S2.** | Temporal smoothing does not fully account for feature convergence | pg. 5-6 |
| **Figure S4.** | Temporal imprecision in the transport plan does not account for feature convergence at recall. | pg. 7-8 |
| **Table S2.** | Linear mixed-effect model estimates and bootstrap confidence intervals. | pg. 8-9 |
| **Table S3.** | Sensitivity of normalized feature convergence to weak recall endpoint signals. | pg. 10 |
| **Table S4.** | Normalized feature separation across all feature-level comparisons. | pg. 10 |
| **Material S1.** | The prompt for GPT-4o API to organize the text | pg. 11 |

**Table S1. Hyperparameter search space and cross-validated selections.**

**(A)** Candidate values evaluated for the Gromov-Wasserstein trade-off parameter $\alpha$ and entropy regularization parameter $\epsilon$. Lower $\alpha$ bias towards more Wasserstein term, higher α bias towards more Gromov term.

| **Parameters** | **Candidate values** |
|---|---|
| $\alpha$, Gromov-Wasserstein trade-off | 0-1 in increments of 0.1 |
| $\varepsilon$, entropy regularization | 0.5-8 in increments of 0.5 |

**(B)** Hyperparameters selected for each held-out participant using leave-one-subject-out (LOSO) cross-validation. Values are reported as ($\alpha$, $\varepsilon$); selection for each participant was based on mean prediction accuracy across the other 15 participants.

| | | | | | | | |
|---|---|---|---|---|---|---|---|
| Sub-01 | Sub-02 | Sub-03 | Sub-04 | Sub-05 | Sub-06 | Sub-07 | Sub-08 |
| 0.3, 8 | 0.3, 8 | 0.3, 8 | 0.3, 8 | 0.2, 8 | 0.3, 8 | 0.3, 8 | 0.3, 8 |
| Sub-09 | Sub-10 | Sub-11 | Sub-12 | Sub-13 | Sub-14 | Sub-15 | Sub-16 |
| 0.3, 8 | 0.3, 8 | 0.4, 8 | 0.3, 8 | 0.4, 8 | 0.4, 8 | 0.3, 8 | 0.3, 8 |

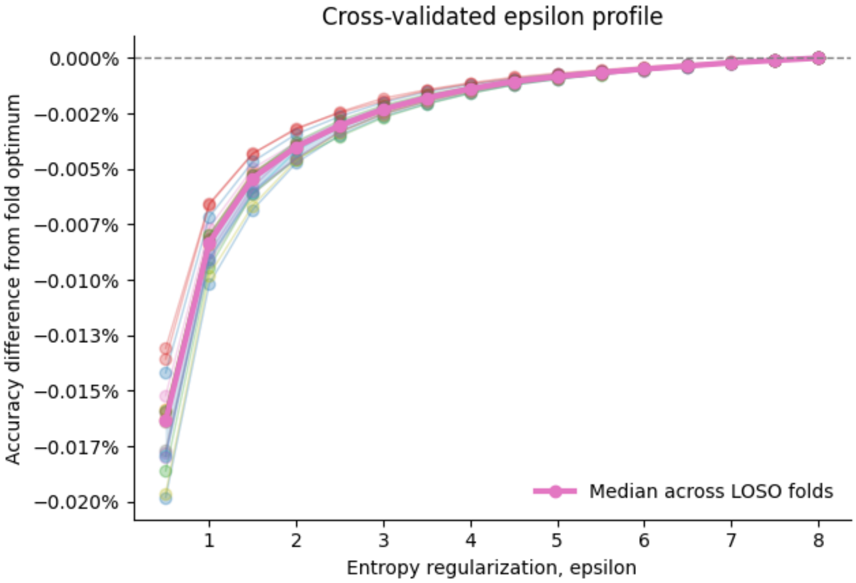


**Figure S1. Cross-validated sensitivity to entropy regularization.** Because every participant's selected ε was at the upper boundary of the grid in Table S1, we tested whether larger values further improved prediction. For each leave-one-subject-out fold, prediction accuracies across all parcels were averaged across the remaining 15 participants for every tested entropy value, after selecting the best-performing $\alpha$ at that value. Accuracy is shown as the difference from the best-performing value within each fold, expressed in percentage points. Thin lines show individual folds, the thick line shows the median across folds, and the shaded band indicates the interquartile range. Validation performance improved with increasing entropy regularization, but the incremental gains diminished markedly at higher values and were negligible near the upper boundary of the tested range. Thus, although ε was selected at the boundary of the grid, performance had effectively plateaued, and the original search range was retained for model fitting throughout the study.

### Supplementary Results S1. OT outperforms diagonal mapping

To confirm OT alignment outperformed simple proportional temporal correspondence, we compared it against a diagonal baseline (each recall TR matched to the encoding TR at the same relative position), using the same feature-prediction pipeline. For each subject and feature level, we summarized the performance using the 90th of normalized rank accuracy across ROIs, then tested the OT versus diagonal difference with a one-sided Wilcoxon signed-rank test across subjects (n = 16).

OT reinstatement accuracy exceeded the diagonal baseline at every feature level. Per-subject differences (OT – diagonal) were reliably positive for low-level visual (median $\Delta = 0.7\%$, $p < .001$; 15/16 subjects above zero), mid-level (median $\Delta = 0.8\%$, $p < .001$; 15/16 subjects), high-level (median $\Delta = 1.2\%$, $p < .001$, 16/16 subjects), and semantic (median $\Delta = 1.2\%$, $p < .001$, 15/16 subjects).

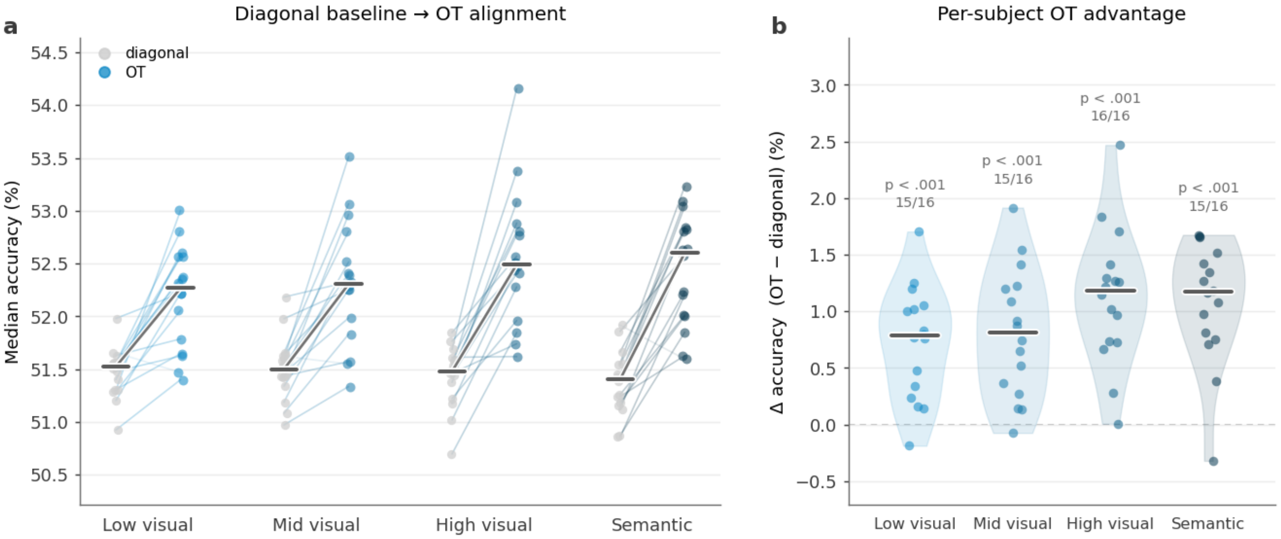


**Figure S2. OT alignment outperforms a diagonal baseline.** Left panel: per-subject reinstatement accuracy for the diagonal baseline (gray) versus OT (color), paired within subject; bold lines indicate group medians. Right panel: per-subject OT vs. diagonal difference by feature level; points are subjects (n = 16), bars are medians, dashed line marks no difference. OT exceeded the baseline at all four feature levels (one-sided Wilcoxon signed-rank).

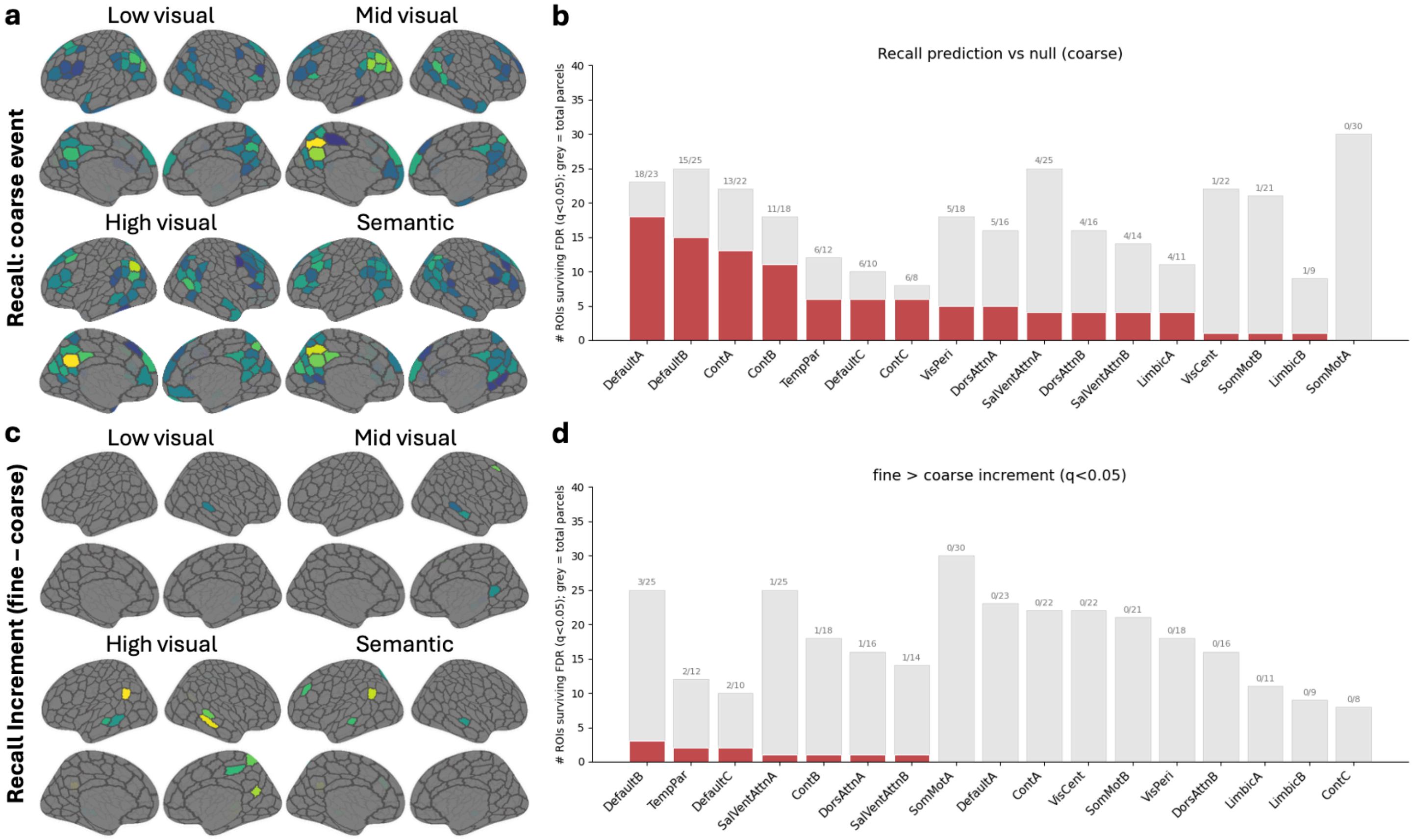


**Figure S3. Temporal granularity comparison for recall. (a)** Recall prediction accuracy at event resolution (coarse). Predicted recall patterns were event-averaged within participant's recall scene spans, so all TRs within an event share a single predicted pattern. Maps show parcels where rank accuracy exceeded a block permutation null (two-sided Wilcoxon across participants, FDR-corrected across parcels within feature level, $q < .05$). **(b)** Network distribution of the parcels in (a). Bars give the number of parcels surviving FDR (red) against the total parcel count in 17-network label (grey). A parcel is counted once regardless of how many feature levels for which it is significant; therefore the map shows a general network reinstatement pattern. **(c)** Recall increment, defined per parcel at rank accuracy at fine (TR) resolution minus rank accuracy at coarse (event) resolution; measures were paired within permutation. **(d)** Network distribution of parcels with a reliable positive increment (one-sided Wilcoxon, same FDR procedure). Because candidate TRs are drawn from the full recall time series, event identity accounts for most of the attainable rank accuracy and the within-event contribution is confined to a narrow margin.

**Supplementary Results S2. Temporal smoothing does not fully account for feature convergence**

**Methods.** Because the recall-side features were obtained by transporting movie features through the fitted transport plan, the blurring induced by it could act like a low-pass filter on the feature timeseries. Specifically, it could attenuate rapidly varying features more than slowly varying ones, so an imprecise plan could reduce feature separation at recall without any change in the represented format. To assess this possibility, we tested whether applying temporal mixing to encoding features could reproduce the reduced feature separation observed during recall; as encoding provides a reference space where movie features have a known one-to-one temporal correspondence with neural activity. Thus, encoding was used to assess the cost of temporal mixing. We examined two forms of temporal mixing, local Gaussian smoothing and non-local mixing based on the empirical OT weights. Both controls retained the original encoding models and neural data while modifying feature time series used for prediction.

For local smoothing, we applied Gaussian kernels with standard deviations of $\sigma = 0, 1, 2, 4, 8, 16, 32$, and 64 TRs (0-96 s), where $\sigma = 0$ represented the original, unsmoothed features. For encoding, we constructed row-normalized Gaussian smoothing matrices, applied them along the movie time and then recomputed predictions at each width. For recall, we reconstructed the transport plan by replacing each row with a delta function at its maximum-weight movie timepoint, then applied Gaussian smoothing over the same range of widths. This allowed us to examine local smoothing without adding further diffusion to the fitted OT plans. To select a width for the encoding-recall comparison, we identified the smallest $\sigma$ at which normalized feature separation under the reconstructed recall plans reached or fell below the separation obtained with the fitted transport plan.

Because local Gaussian smoothing does not capture mixing between temporally distant movie moments, we also constructed a non-local control using the empirical weights from the OT plans fitted for viewing-recall alignment. For each time point in the test encoding session, we selected the row of the fitted OT plan whose peak weight was the closest to that movie time point. We then shifted the row to align its peak with the target time point to minimize the displacement while preserving the relative temporal offsets and weights of the remaining contributions. Weights extending beyond the combined timeline of both encoding sessions were discarded, and each row was renormalized to sum to one. Now, the temporal alignment between encoding features and neural activity were maintained with preserved temporal mixing structure of the empirical OT plan. To accommodate contributions from either movie session, features from both sessions were projected into the PCA space fitted on the data from the session that the encoding model was trained. The resulting mixing matrix was applied to all four feature levels before recomputing predictions using the previously fitted encoding models.

For both controls, predictive performance was evaluated using rank accuracy within the same fixed cortical mask used in the primary analyses. Normalized feature separation was defined as the semantic-minus-low-visual difference in above-chance rank accuracy divided by the mean above-chance accuracy of these two feature levels. Encoding estimates were averaged across the two training directions within each participant. Encoding and recall separation were compared using two-sided Wilcoxon signed-rank tests across participants. For visualization, group-level estimates are reported as medians across participants, with error bars indicating the interquartile range.

**Results.** Gaussian smoothing progressively reduced above-chance rank accuracy across feature levels (Fig. S4b). However, the normalized feature separation remained largely preserved across the moderate smoothing range (Fig. S4a), suggesting that the temporal smoothing did not by itself reproduce the convergence we observed during recall (Fig. S4c). At extreme smoothing widths, prediction approached chance.

For the reconstructed recall plans, normalized feature separation fell to the level obtained with the OT plan at $\sigma = 4$ TRs (Fig. S4c). We therefore used this width as the encoding side comparison point, as it is the amount of imprecision that reproduces the separation the reconstructed plan yields. At this width, encoding feature separation remained four times that of recall (0.796 vs. 0.200; 14/16 participants, two-sided $p < .001$), indicating that local Gaussian smoothing did not reproduce the observed convergence.

The non-local control introduced wide temporal mixing while preserving nearly all the original OT weights after shifting (median proportion retained = 100%). Across participants and encoding prediction directions, a median of 79.4% of the transport mass was assigned to movie time points more than 100 TRs from the target point, indicating a clear non-local mixing pattern. This mixing reduced predictive performance, with the median endpoint accuracy above chance decreasing from 0.014 to 0.010 (Fig. S4f). Despite this reduction, normalized feature separation instead increased from 0.788 during original encoding to 1.087 under non-local mixing (13 of 16 participants, two-sided $p = .034$), while remaining greater than 0.200 during fitted-OT recall (15 of 16 participants, two-sided $p < .001$; Fig. S4e). Thus, the non-local mixing did not reproduce the feature convergence observed during recall even though it weakened the encoding predictive performance.

Together, these controls showed that neither local Gaussian smoothing nor the non-local mixing reproduced the feature convergence effect during recall. These findings argue against transport-induced temporal mixing as an explanation for feature convergence and supports the notion that recall reorganizes the relative representation of different feature levels beyond the effects of temporal alignment imprecision.

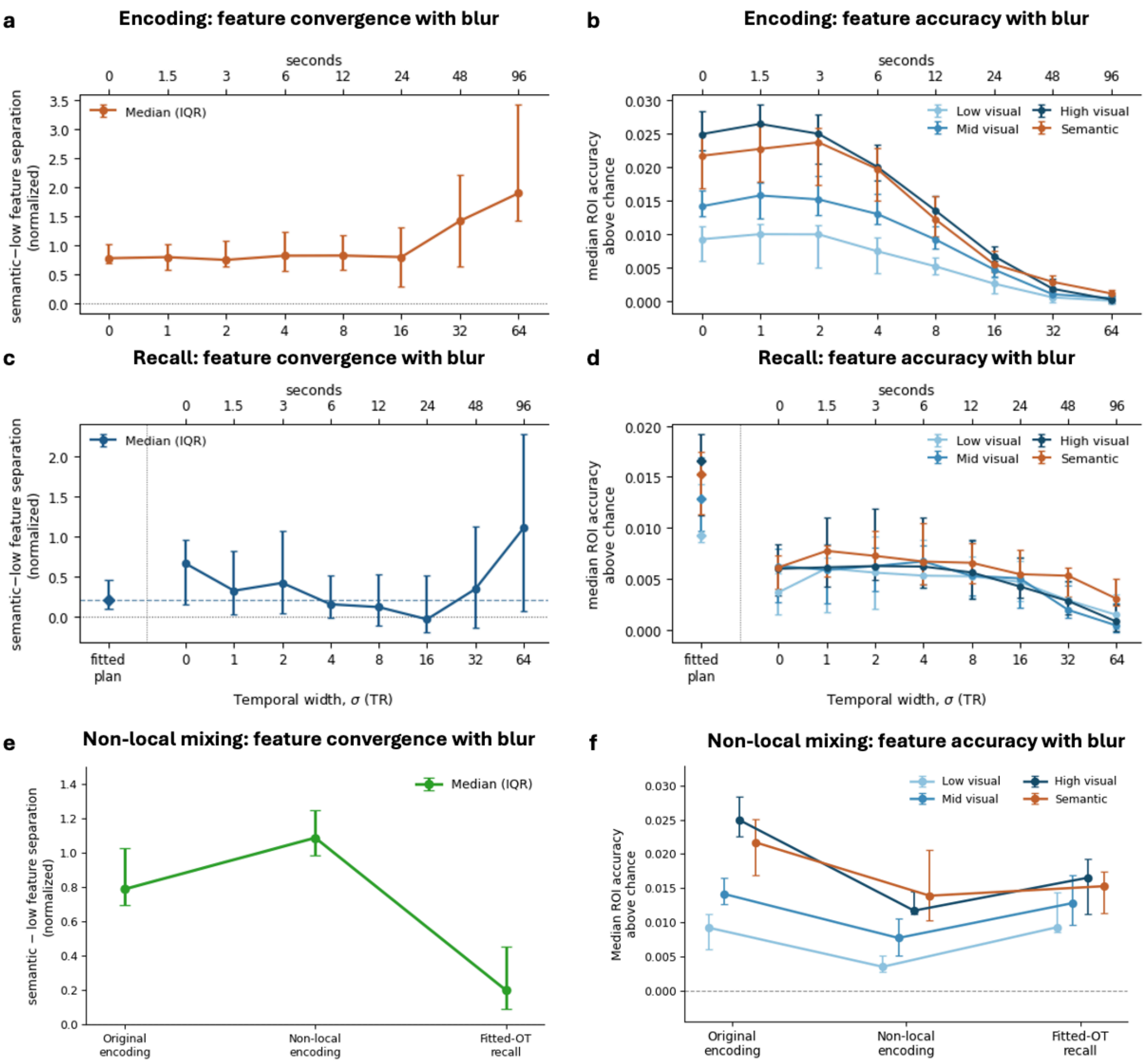


**Figure S4. Temporal imprecision in the transport plan does not account for feature convergence at recall.** Points are medians across participants, and error bars indicate the interquartile range (25th-75th percentiles). Analyses were restricted to the fixed recall-defined union mask (parcels showing evidence for at least one feature level at uncorrected $p < .001$). Feature-level accuracy is the median rank accuracy across parcels within this mask, expressed relative to chance (0.5). Normalized semantic-low level separation is the difference between semantic and low-level above-chance accuracy divided by their mean, making the measure invariant to uniform scaling of the two feature signals. Lower values indicate greater convergence. Blur is Gaussian with width σ, shown in TRs (lower axis) and seconds (upper axis). **(a)** Encoding: normalized feature separation as a function of blur width. Because encoding timing is veridical, blurring it by σ distributes each timepoint over its neighbours. Values are averaged within participant across the two cross-validation directions before separation is computed. **(b)** Encoding: above-chance accuracy for each feature level across the same sweep. **(c)** Recall: normalized feature separation. The leftmost point (diamond, separated by the vertical line) is the fitted transport plan, and the dashed horizontal line marks its median. Remaining points are plans reconstructed to have width exactly σ: each row of the fitted plan is replaced by a delta function at its peak movie timepoint and re-blurred by σ. **(d)** Recall: above-chance accuracy for each feature level, conventions as in (c). **(e)** Normalized feature separation during original encoding, encoding with non-local mixing, and fitted-OT recall. The non-local control applied recall-derived OT weight distributions to encoding features, selecting and shifting each distribution to align its peak with the target encoding time point while retaining contributions from temporally distant movie moments. **(f)** Above-chance predictive accuracy for all four feature levels under the same three conditions as in (e). Non-local mixing reduced predictive accuracy but did not reproduce the reduction in normalized feature separation observed during recall.

**Supplementary Table S2. Linear mixed-effect model estimates and bootstrap confidence intervals.** Rank accuracy in percentage points (chance = 50). Feature level sum-to-zero coded; gradient mean-centered across parcels. Contrasts evaluated at the mean gradient position.

**(a) Omnibus test.**

| Term | df1 | df2 | F | p |
|---|---|---|---|---|
| State | 1 | 15.0 | 96.71 | < 0.001 |
| Feature | 3 | 38032 | 492.48 | < 0.001 |
| Gradient | 1 | 298.0 | 0.50 | 0.480 |
| State × Feature | 3 | 38032 | 291.89 | < 0.001 |
| State × Gradient | 1 | 38032 | 1453.18 | < 0.001 |
| Feature × Gradient | 3 | 38032 | 6.43 | < 0.001 |
| State × Feature × Gradient | 3 | 38032 | 16.91 | < 0.001 |

**(b) Pairwise feature-level separations within state** (estimated marginal means, Wald 95% CI, no multiplicity adjustment). * denotes 95% confidence interval (CI) excludes 0.

| Contrast | Encoding | 95% CI | Recall | 95% CI |
|---|---|---|---|---|
| mid – low | 0.716 | [0.652, 0.779]* | 0.057 | [-0.006, 0.120] |
| high – low | 1.510 | [1.446, 1.573]* | 0.181 | [0.117, 0.244]* |
| semantic - low | 1.048 | [0.984, 1.111]* | 0.169 | [0.105, 0.232]* |
| high – mid | 0.794 | [0.730, 0.857]* | 0.124 | [0.060, 0.187]* |
| semantic – mid | 0.332 | [0.269, 0.395]* | 0.112 | [0.048, 0.175]* |
| semantic – high | -0.462 | [-0.525, -0.398]* | -0.012 | [-0.075, 0.052] |

**(c) Encoding to recall change in prediction accuracy, by feature level**

| Feature level | Recall – encoding | 95% CI |
|---|---|---|
| low visual | -0.365 | [-0.587, -0.142] |
| mid visual | -1.024 | [-1.246, -0.801] |

| | | |
|---|---|---|
| high visual | -1.694 | [-1.916, -1.471] |
| semantic | -1.244 | [-1.466, -1.021] |

**(d) Gradient slopes**

| Feature level | Encoding | 95% CI | Recall | 95% CI | Change |
|---|---|---|---|---|---|
| low visual | -0.065 | [-0.089, -0.041] | 0.068 | [0.044, 0.092] | 0.133 [0.117, 0.149] |
| mid visual | -0.068 | [-0.092, -0.044] | 0.090 | [0.066, 0.114] | 0.158 [0.142, 0.174] |
| high visual | -0.106 | [-0.130, -0.082] | 0.101 | [0.077, 0.125] | 0.207 [0.191, 0.224] |
| semantic | -0.047 | [-0.071, -0.023] | 0.089 | [0.065, 0.113] | 0.136 [0.120, 0.153] |
| averaged | -0.072 | [-0.093, -0.050] | 0.087 | [0.065, 0.109] | 0.159 [0.124, 0.190] |

**(e) Contrasts with subject-level bootstrap CIs** (500 resamples, subjects resampled with replacement)

| Quantity | Contrast | Estimate | 95% bootstrap CI |
|---|---|---|---|
| Gradient shift | slope change, averaged over levels | 0.158 | [0.124, 0.190] |
| Feature convergence | (semantic-low at recall) – (semantic-low at encoding) | -0.873 | [-1.068, -0.695] |
| Three-way, high vs low | slope change difference | 0.075 | [0.047, 0.100] |
| Three-way, mid vs low | slope change difference | 0.025 | [0.000, 0.048] |
| Three-way, semantic vs low | slope change difference | 0.005 | [-0.029, 0.039] |
| Overall level change | mean accuracy, recall – encoding | -1.071 | [-1.272, -0.859] |

**Supplementary Table S3. Sensitivity of normalized feature convergence to weak recall endpoint signals.** Normalized feature separation was defined as $S = (SM - LV)/[(SM + LV)/2]$, where $SM$ and $LV$ are semantic- and low-level-feature rank accuracies after subtracting chance (0.5). Because this measure expresses the difference between these two endpoint features relative to their mean above-chance prediction strength, estimates may be less stable when both endpoint signals approach chance. We therefore repeated the encoding-recall comparison by progressively excluding participants with weak recall endpoint signals, using thresholds of 0.10, 0.25, and 0.50 percentage points above chance. Normalized feature separation remained substantially lower during recall across all thresholds, with median recall-minus-encoding differences ranging from -0.655 to -0.674. P values are from one-sided Wilcoxon signed-rank tests testing lower normalized separation during recall. The stability of the effect across thresholds indicates that feature convergence was not driven by participants whose normalized separation was estimated from weak recall signals.

| **Minimum recall endpoint signal** | **N** | **Median encoding S** | **Median recall S** | **Median recall – encoding** | **p-value** |
|---|---|---|---|---|---|
| No exclusion | 16 | 0.788 | 0.200 | -0.552 | < .001 |
| > .0025 | 16 | 0.788 | 0.200 | -0.552 | < .001 |
| > .0050 | 15 | 0.761 | 0.194 | -0.598 | < .001 |
| > .0075 | 13 | 0.761 | 0.206 | -0.507 | < .001 |

**Supplementary Table S4. Normalized feature separation across all feature-level comparisons.** To determine whether the reduction in feature separation was specific to the low-level visual-semantic endpoints used in the main analysis, we computed normalized separation for each of the six pairwise comparisons among low-, mid-, high-level visual, and semantic features. For each pair, separation (S) was defined as the absolute difference in above-chance rank accuracy divided by the mean above-chance rank accuracy of the two feature levels, matching the normalization used for the endpoint analysis in main text. Values are participant medians. Recall-minus-encoding values below zero indicate reduced feature separation during recall. One-sided Wilcoxon signed-rank tests assessed whether normalized separation was lower during recall than encoding, with Holm correction across the six comparisons.

| **Feature-level comparison** | **Median encoding S** | **Median recall S** | **Median recall – encoding** | **p-Holm** |
|---|---|---|---|---|
| low – mid | 0.728 | 0.210 | -0.431 | .027 |
| low – high | 1.076 | 0.232 | -0.687 | .013 |
| low – semantic | 0.863 | 0.342 | -0.541 | .018 |
| mid – high | 0.498 | 0.119 | -0.252 | .005 |
| mid – semantic | 0.334 | 0.253 | -0.054 | .433 |
| high – semantic | 0.177 | 0.170 | -0.038 | .433 |

**Supplementary Material S1. The prompt for GPT-4o API to organize the text**

GPT-4o API was used as a preprocessing step to combine the encoding scene context and scene annotation available from Chen et al. (2017) into a single sentence. The model was instructed to preserve only the information provided in the original inputs and not to introduce additional details with temperature set as 0. The following prompt was used:

| **Prompt** |
| --- |
| Here is context about the scene:<br>[scene context]<br><br>And here is scene description:<br>[scene details]<br><br>Please rewrite only the information given above as a single sentence that begins with the context and then describes the scene. Do not add any new details, adjectives, sensory descriptions, or labels. |

One example of the rewriting process is:

| | |
| --- | --- |
| **Context** | Soldiers in Afghanistan |
| **Scene details** | Gunfire by a soldier along a wall made of stacked stones |
| **Rewritten sentence** | In Afghanistan, a soldier fires a gun along a wall made of stacked stones. |
